# The oral microbiome of King Richard III of England

**DOI:** 10.1101/2025.09.21.677585

**Authors:** Irina M. Velsko, Alexander Hübner, Zandra Fagernäs, James Fellows Yates, Allison E. Mann, Courtney Hofman, Andrew T. Ozga, Cecil M. Lewis, Camilla Speller, Sarah Fiddyment, Michael Francken, Joachim Wahl, Johannes Krause, Anita Radini, Turi King, Christina Warinner

**Author notes:** **Corresponding Authors** Irina Velsko, Turi King, Christina Warinner.

## Abstract

Metagenomic investigation of archaeological dental calculus has provided insights into the changing oral health, disease, and diet of past human societies, but little is known about the oral microbiota of exceptionally high-status individuals, whose diet and lifestyle sharply differed from the general population. Here we analyze the dental calculus metagenome of King Richard III of England (1452-1485) and compare it to new and previously published dental calculus metagenomes from predominantly non-elite contexts in England, Ireland, the Netherlands, and Germany spanning the Neolithic to the present. Deep sequencing and *de novo* assembly enabled the investigation of metagenome-assembled genomes (MAGs) from periopathogens within the genus *Tannerella*. DNA preservation within the dental calculus of King Richard III was found to be exceptional, yielding an extraordinarily well-preserved oral microbiota. Oral microbiome species diversity fell within the range previously observed among other northern European populations over the past 7,000 years, suggesting that a royal lifestyle and a rich diet did not substantially shift his oral microbiome composition. Reconstructed *Tannerella* genomes contained many virulence factors found today among periopathogens, including *Tannerella forsythia*. Insufficient plant and animal DNA was recovered to investigate diet, suggesting that dental calculus may not be a sufficient source of dietary DNA for dietary reconstruction, even when well-preserved. The dental calculus of King Richard III has produced one of the richest and best preserved ancient oral metagenomes studied to date and contributes to understanding the ecology and evolution of the human oral microbiome.

## Introduction

In 2012, the skeletal remains of King Richard III of England were discovered beneath a car park on the grounds of the former Grey Friars church in the city of Leicester, UK [1]. Subsequent osteological [1–3] and genetic and statistical [4] analyses of the evidence confirmed the identity of the remains of the long lost king. Richard III became king of England in 1483 during a period of civil conflict between the Plantagenet Houses of York and Lancaster later known as the Wars of the Roses (1455-1487). Remembered as a usurper, Richard III was accused by chronicler John Rous [5] and Sir Thomas More [6] of plotting the murder of his nephews to seize the throne. Crowned king at age 30, he reigned for two tumultuous years and was killed at the Battle of Bosworth in 1485, after which his body was hastily buried at the Grey Friars Church in Leicester. His life and brief reign became the subject of an eponymous tragic play by William Shakespeare in 1593 in which Richard III was portrayed as a ruthless villain [7]. Although this literary legacy has made him one of England’s most well-known medieval kings, relatively little is known about his life other than its political outline, and the details of his rise to power following his brother’s death remain sharply disputed [8]. Archaeology provides an opportunity to augment his scant historical record.

The discovery of Richard III’s remains has provided an unusual chance to not only confirm specific historical details of his death but also to investigate less well-understood aspects of his life. Analysis of his remains has confirmed his death in battle, as his skeleton bore the marks of at least eleven perimortem wounds, including two likely fatal wounds to the back of the head and several humiliation injuries that may have been inflicted after he was stripped of armor [2]. Historical accounts of a hasty burial have also been supported by the absence of evidence for a coffin, or even shrouding, with his corpse having been placed within a cramped grave, possibly with his hands bound [1]. Accounts of his uneven shoulder height are supported by the identification of idiopathic adolescent-onset scoliosis in his skeleton [3], which likely began during his training as a knight in Yorkshire. Well-preserved DNA within his skeleton has confirmed his sex as male, and mitochondrial genome sequencing has conclusively linked him to two female-line relatives [4]. Isotopic analysis of his enamel bioapatite (Sr, Pb, O) and collagen from dentine and bone (O, C, N) has provided further insights into aspects of his life history [9]. His oxygen and strontium isotopic values are consistent with his childhood movements between eastern and western England, and his carbon and nitrogen isotopic values reflect a changing diet in childhood, followed by a likely increase in meat and high trophic level fish consumption from his teenage years onwards, and later a likely further increase in the regular consumption of luxury foods, such as wine, fish, and wildfowl, after being crowned king.

The excellent biomolecular preservation of the remains of Richard III suggests that an analysis of the DNA present in his dental calculus might yield further insights into his life, and specifically on his oral health and diet. Dental calculus, a mineralized form of dental plaque also known as tooth tartar, forms incrementally during life, entrapping the oral microbiota and serving as a long-term biomolecular archive of the oral microbiome [10,11]. Food biomolecules may also become entrapped in dental calculus and can further aid in dietary reconstruction [12,13]. However, while microbial metagenomics has been widely applied to reconstruct the microbial structure and diversity of the oral microbiome through time [14,15] and to track the prevalence and virulence of periodontal pathogens [16–19], the use of dental calculus metagenomics to study diet has been more limited [20], and many challenges remain for authentication of DNA from dietary sources [21].

Studies of the impact of diet on oral microbiome composition in living populations suggest that although individual microbial species may differ in abundance between populations with different diets, the overall species composition remains consistent across a variety of diets [14,22–25]. To date, studies of ancient dental calculus have likewise found relatively little difference in the microbial composition of calculus from populations practicing different subsistence strategies [20,26–28]. This leaves us with an unclear understanding of whether and how social status may have historically impacted oral microbiota. To date, the majority of published metagenomic data from ancient dental calculus come from individuals who were not of high social status [29–31] or whose social status is unknown. Analyzing the dental calculus metagenome of King Richard III offers the opportunity to investigate the oral microbiota and diet of a high-status individual for which there are both historical records and archaeological data for comparison.

Here, we metagenomically characterize dental calculus from King Richard III and assess the presence of dietary DNA, as well as compare the microbial species profile to new and published dental calculus of predominantly non-elite individuals from locations in northern Europe spanning the last 7,000 years. We found that the dental calculus of Richard III contains an extraordinary amount of well-preserved DNA, and that the microbial species composition is indistinguishable from that of other historic-era calculus from across northern Europe, suggesting that a royal lifestyle and a rich diet did not substantially alter his oral microbiota. We reconstructed metagenomically-assembled genomes (MAGs) within the genus *Tannerella*, and found that the periopathogen *Tannerella forsythia* of Richard III falls within the known diversity and virulence of other modern and ancient *T. forsythia* in European populations. Putative dietary sequences could not be authenticated. Although deeper sequencing may provide additional dietary DNA sequences, the very low proportion of putative dietary sequences suggests that not all dental calculus may be a sufficient source of dietary DNA for dietary reconstruction, even when exceptionally well-preserved.

## Results

### Exceptional DNA recovery from the dental calculus of Richard III

We extracted DNA from four calculus samples collected from three teeth of Richard III (Table S1), and during laboratory processing we observed very high recovery of DNA, ranging from 122.4-506.7 ng/mg (Table 1). We compared this DNA recovery to 143 other archaeological dental calculus samples previously measured in our laboratories (Table S2), which ranged from 0.1 ng/mg to 437 ng/mg, with a median of 36 ng/mg (Figure 1A). We found that the amount of DNA recovered from the calculus of King Richard III is among the highest we have ever measured from an archaeological context, and is comparable to that at other recent archaeological sites with exceptional preservation in northern Europe (Dalheim, Tickhill, Middenbeemster) and Oceania (Rapa Nui). Such high DNA concentrations are similar to those measured from modern calculus (83.4-346 ng/mg) and suggest an excellent state of DNA preservation. To take advantage of this preservation and allow *de novo* genome assembly, we sequenced each dental calculus library from Richard III to a high depth totaling more than 400 million sequences (Figure 1B). To facilitate comparative analysis, we also performed DNA extraction and deep sequencing of dental calculus from three individuals from the Neolithic Linearbandkeramik (LBK, 8th millennium BP) site of Stuttgart-Mühlhausen (Table S1) in Baden-Württemberg, Germany [32], and performed deeper sequencing of dental calculus libraries (Table S1) from the well-preserved G12 individual at the site of Dalheim, Germany (medieval, 750-1000 BP) [12]. We then selected and downloaded published dental calculus metagenomic datasets within the AncientMetagenomeDir (61) dating to 200-1000 BP from England, Ireland, the Netherlands, and Germany (Table S3) for comparative microbiome analysis.

**Table 1.** DNA extraction results.

| Sample ID | Tooth | Calculus analyzed (mg) | Elution vol (μl) | DNA yield (ng/μl) | Normalized DNA yield (ng/mg) |
| --- | --- | --- | --- | --- | --- |
| R3-1 | LP<sub>1</sub> | 3.6 | 60 | 30.4 | 506.7 |
| R3-2A | LC<sub>1</sub> | 1.0 | 60 | 2.0 | 122.4 |
| R3-2B | LC<sub>1</sub> | 2.3 | 60 | 6.3 | 163.3 |
| R3-3 | RP<sub>2</sub> | 4.0 | 60 | 25.8 | 387.0 |
| R3-AncBL1 | - | - | 60 | * | - |
| R3-AncBL2 | - | - | 60 | 0.2 | - |

Notes: R3, Richard III; \*, below detection.

**Figure 1.**
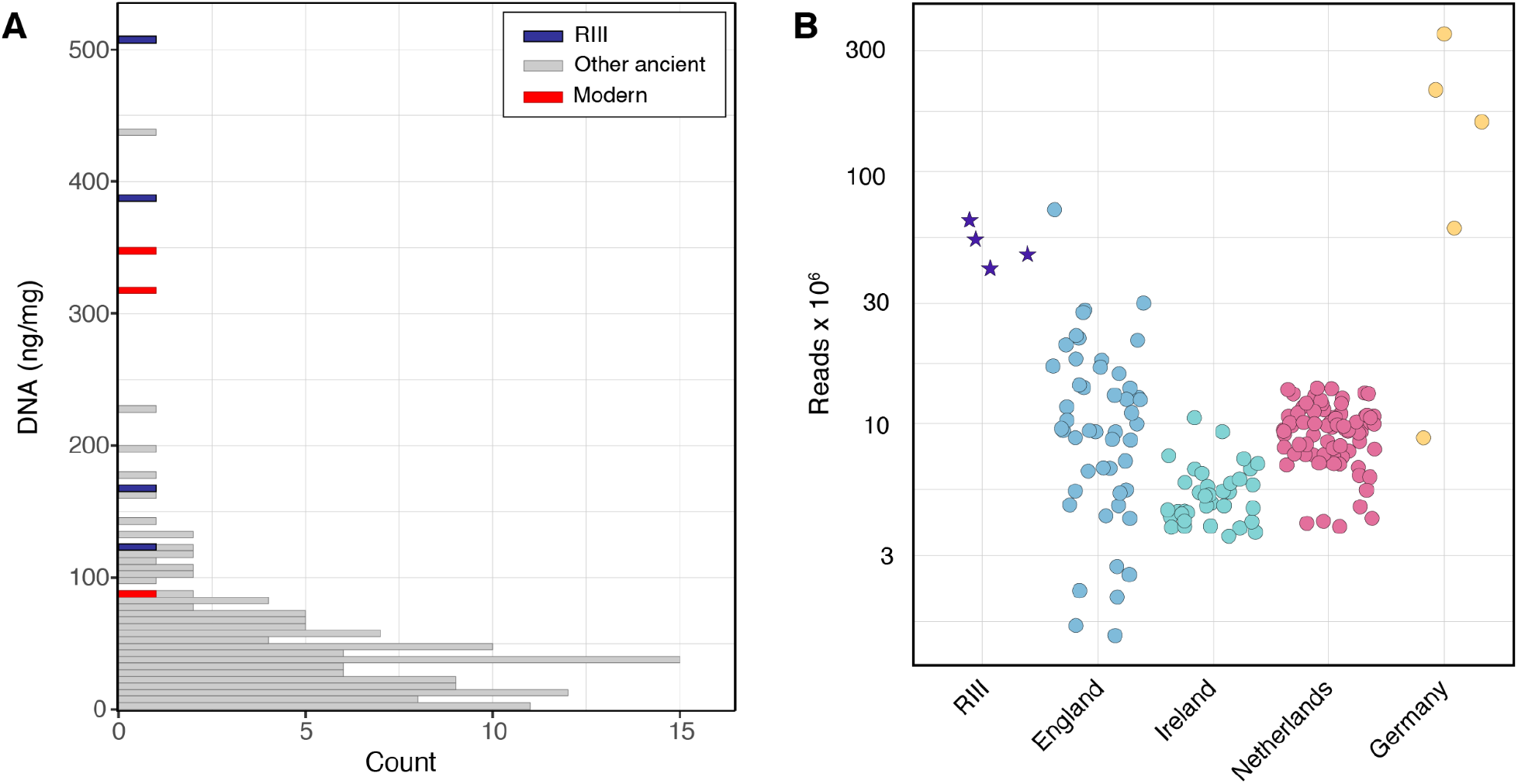
Quantification of extracted DNA and sequenced reads. **A**. Normalized DNA recovery reported in ng DNA / mg dental calculus, with the dental calculus of Richard III (RIII) indicated in purple, other ancient dental calculus in gray, and modern dental calculus in red. **B**. The number of DNA sequences per individual analyzed in this study (except the four libraries from Richard III which are reported separately), including both newly generated and published sequencing data after read processing and quality filtering; note use of log scale.

### Well-preserved oral microbiota but no confirmed dietary DNA

Taxonomic classification of newly sequenced dental calculus from Richard III, Stuttgart-Mühlhausen, and the medieval site of Dalheim with kraken2 [33,34] using the GTDB [35] indicated that all samples were exceptionally well-preserved, with > 90% of DNA sequences attributed to an oral source (dental calculus or dental plaque) based on SourceTracker analysis (Figure 2; Table S4). Additional preservation assessment using cuperdec [14] confirmed a high proportion of oral taxa in these samples, as well as in the comparative dental calculus datasets (Figure S1).

**Figure 2.**
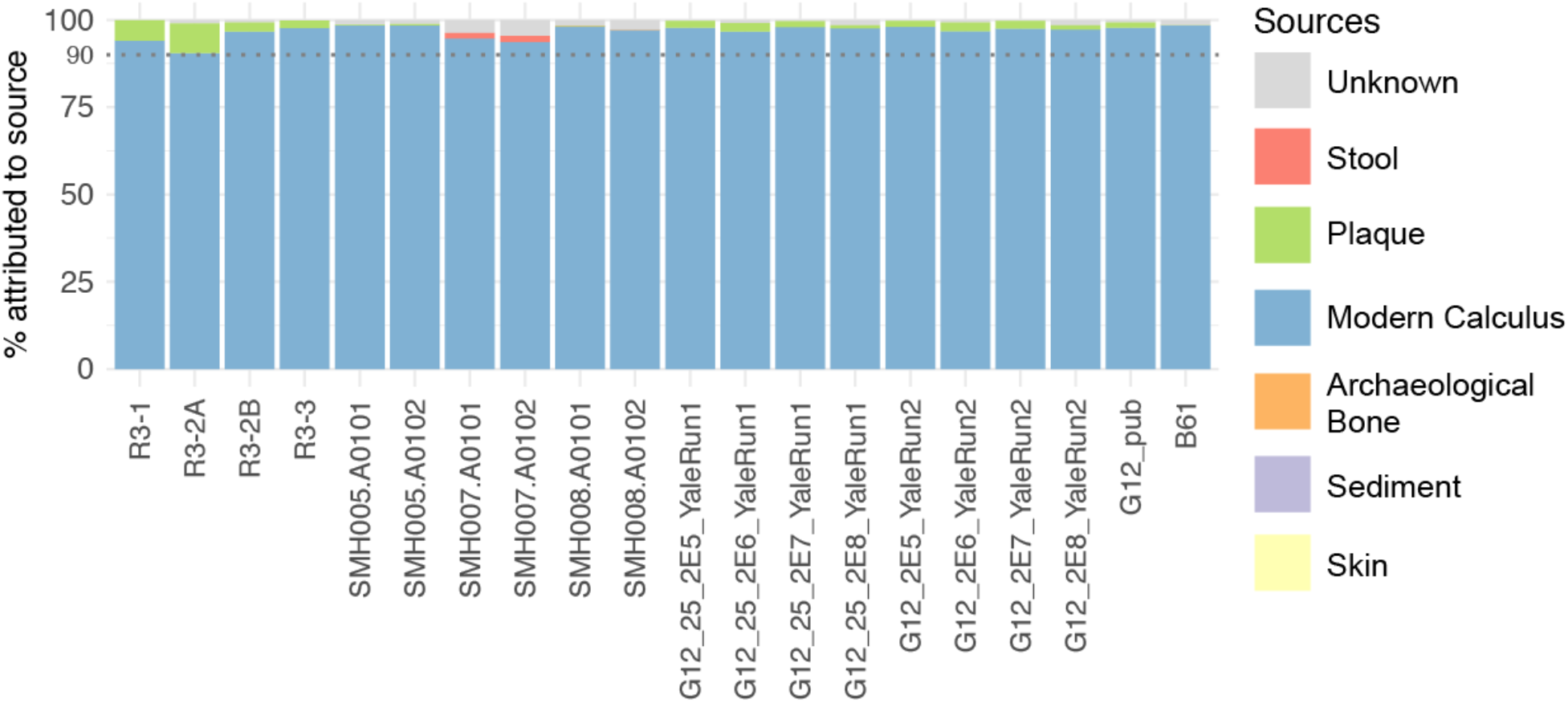
SourceTracker analysis of dental calculus preservation. All newly sequenced DNA libraries in this study exhibit a high preservation of oral microbial species with minimal contamination from other sources. Published data for sample G12 (G12_pub) and B61, from the same site, are shown for comparison. Dotted line indicates 90%.

Within the dental calculus of Richard III, however, no dietary sequences could be authenticated or confirmed. No plant sequences were identified following taxonomic classification with MALT [36,37] using the NCBI nt database, and non-human animal sequences were limited to two species known to have contaminated and unreliable reference genomes [21]: *Cyprinus carpio* (common carp, 4,112 reads) and *Erinaceus europaeus* (European hedgehog, 42,683 reads) (Table S5). These two taxa were also identified in negative and environmental controls, further supporting their identification as false positives. Modern dental plaque likewise contained few dietary reads, with the largest number of reads assigned to contaminated genomes (*Cyprinus carpio*), model organisms (e.g., *Danio rerio*), and wheat (*Triticum aestivum*) (Table S5). This suggests that dietary DNA incorporation and preservation within oral biofilms may be limited.

In contrast to dietary DNA, recovery of DNA from the oral microbiota was excellent. We identified nearly four hundred microbial species in the dental calculus of Richard III, which is similar to the number of species detected in other well-preserved archaeological dental calculus samples from England, Ireland, the Netherlands, and Germany (Figure 3A). Species evenness as measured by the Shannon index was similar across the dental calculus data sets (Figure 3B). Analysis of between-sample diversity places the dental calculus of Richard III within the known microbial diversity of northern Europe (Figure 3C), indicating that the oral microbial profile of the king was not notably different from that of other individuals living in a broadly similar geographic region and time period. The main difference between the king’s calculus and other archaeological dental calculus from northern Europe appears to be primarily the amount of microbial DNA preserved and not which microbial species were present or how abundant they were.

**Figure 3.**
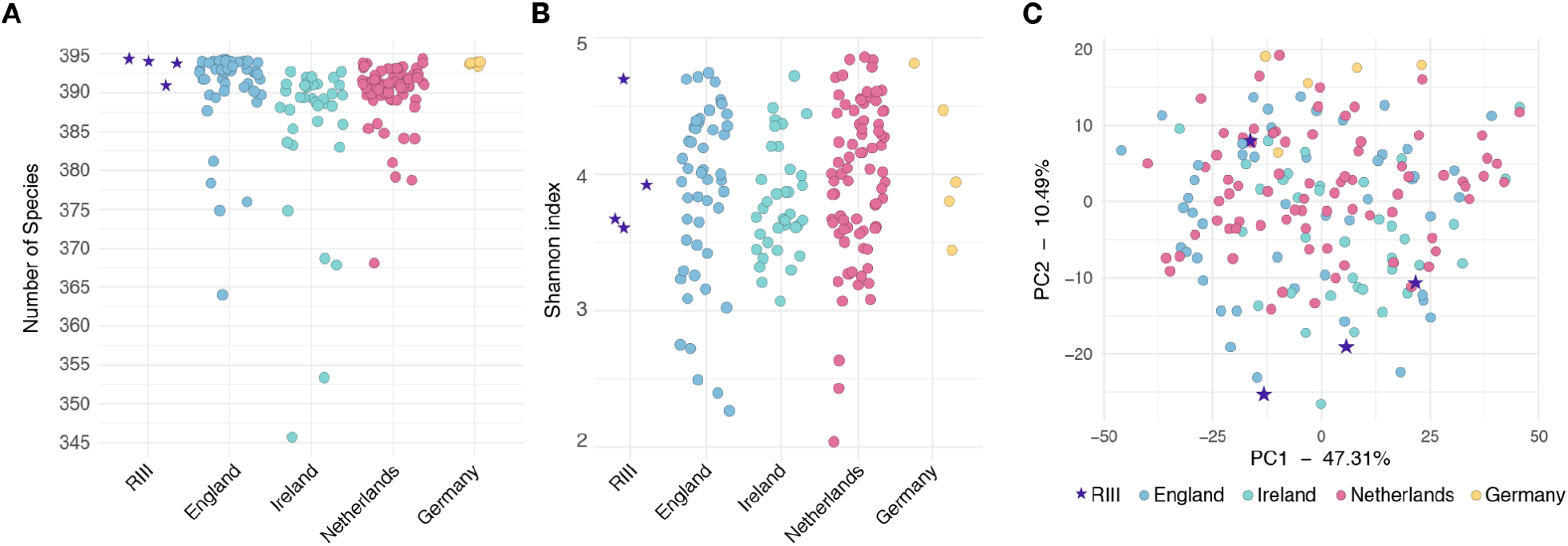
Microbiome metagenomic diversity analysis comparing calculus from King Richard III to other archaeological individuals in northern Europe. **A**. Number of microbial species detected. **B**. Shannon index of species evenness. **C**. Principal component analysis (PCA) of between-sample microbial diversity.

### *Phylogenetic and virulence factor diversity of* Tannerella forsythia

One species found in high abundance and with good preservation in the dental calculus of Richard III was *Tannerella forsythia. T. forsythia* is a common oral species that is today associated with periodontal disease [38], but which was historically highly abundant in calculus on teeth with no evidence of disease [30]. We performed *de novo* assembly and binning of the dental calculus metagenomes, and recovered one metagenome-assembled genome (MAG) of *T. forsythia* from Richard III, four from the Radcliffe Infirmary in Oxford, England, and three from Dutch site of Middenbeemster. We also recovered four genomes of an unnamed *Tannerella* species from the Radcliffe Infirmary, and one genome each from the sites of Stuttgart-Mühlhausen and Dalheim in Germany. All fourteen *Tannerella* genomes meet the minimum quality thresholds for medium and high quality MAGs [39] used for modern-day genome reconstruction (Table S6, Table S7).

To confirm an ancient origin for the reconstructed *Tannerella* genomes, we mapped the reads from each individual (except the four libraries from Richard III which were each analyzed separately) that produced a *Tannerella* MAG against the *T. forsythia* reference genome and assessed the 5’ C-to-T damage patterns (Figure 4A). DNA sequences from Richard III and the Radcliffe Infirmary produced expected age-related damage patterns, indicating that the *Tannerella* sequences in these samples originate from an ancient source [40]. Libraries from the Dutch site of Middenbeemster were prepared with partial-USER treatment and showed an expected ancient DNA damage pattern of reduced damage only on the first base [41]. Libraries from the German sites were prepared with full-USER treatment (Stuttgart-Mühlhausen) or a proofreading enzyme (Dalheim) and consequently show no DNA damage [42,43]. Notably, only the samples from the Netherlands show a baseline damage frequency of 0, while the others range between 0.025 and 0.075, suggesting that other *Tannerella* species may be present in these samples. As only a single MAG was recovered from each assembly, it is likely that additional *Tannerella* species are present at lower abundance, and hence did not produce a MAG during assembly due to insufficient read counts. At present, four *Tannerella* species are recorded as inhabiting the human oral cavity in the Human Oral Microbiome Database (eHOMD) v.4 [44,45], but reference genomes are available for only two: *T. forsythia* and *T. serpentiformis*. Further work is needed to investigate the diversity of this genus to better understand its ecology in ancient dental calculus.

**Figure 4.**
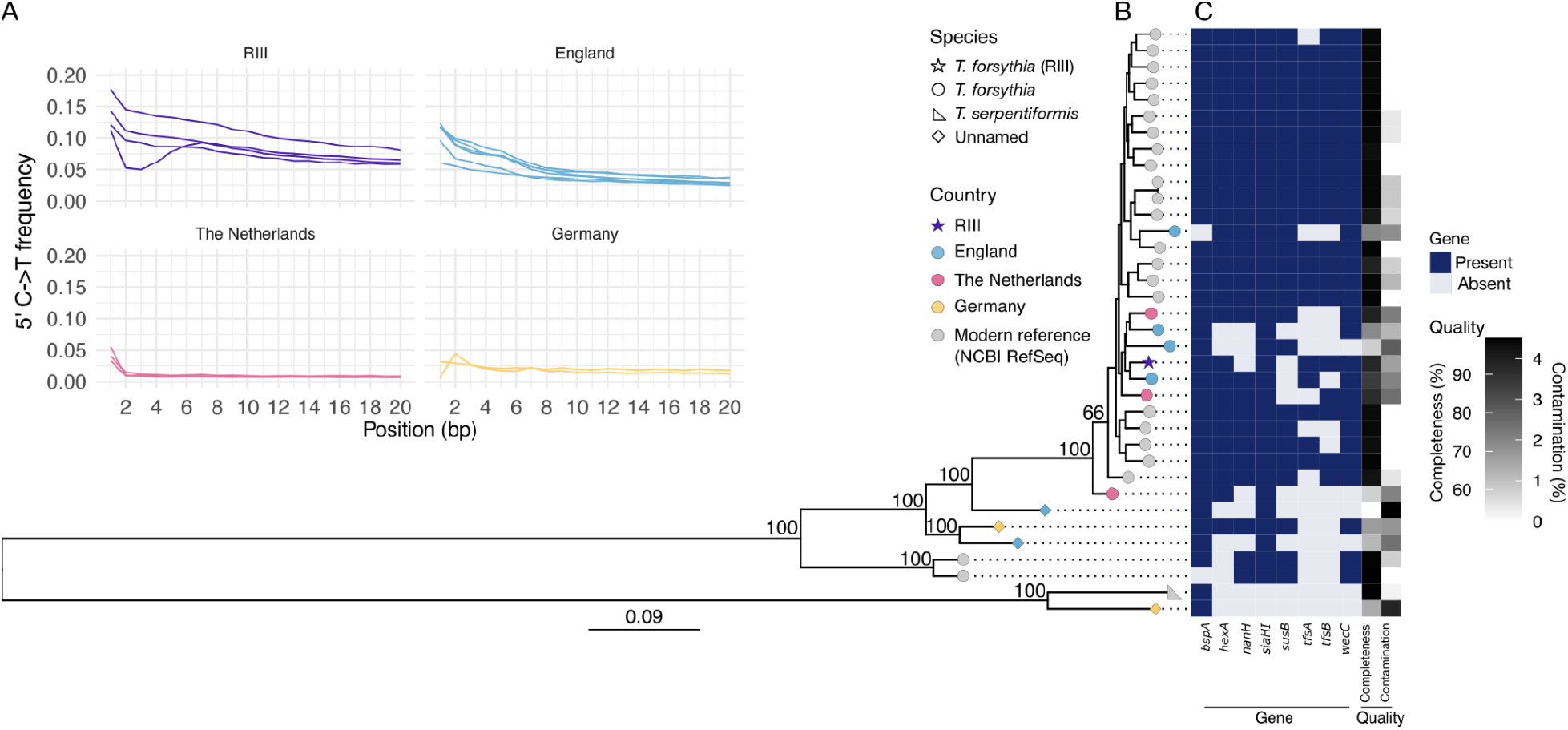
Analysis of ancient *Tannerella forsythia* phylogeny and virulence. **A**. 5’ C-to-T damage patterns for each dental calculus sample (each dental calculus library for Richard III) mapped against the *T. forsythia* reference genome, grouped by country of origin. Richard III and other samples from England show expected damage patterns, while samples from the Netherlands were partial-USER-treated and show correspondingly low levels of damage on the first base. Damage was not observed for the German samples because they were processed using full-USER treatment or a proofreading enzyme during library construction. **B**. Phylophlan single-marker gene phylogeny of reference and ancient *de novo* reconstructed *Tannerella forsythia* genomes, mid-point rooted. RAxML bootstrap values of deep nodes are shown. **C**. Presence/absence matrix of virulence factors for *T. forsythia*, as well as genome completeness and contamination, aligned with the tree tips in panel B. Genome labels are provided in Figure S2.

Phylogenetic analysis of the *de novo* assembled genomes, along with reference genomes of *T. forsythia* from NCBI and the outgroup genome of *T. serpentiformis*, placed the ancient *T. forsythi*a genomes from Richard III, England, and the Netherlands in a clade along with most of the reference genomes (Figure 4B). Two reference genomes and three of the unnamed *Tannerella* genomes (two from England and one genome from Neolithic Germany) fall outside of the main *T. forsythi*a clade, suggesting these belong to different strains, and possibly different but currently unnamed species. Patristic distance calculations support this designation, as values within the *T. forsythia* clade are 0.053 when excluding these genomes, which is at the boundary of species-level cut-offs (0.05) [46], while the average patristic distance within the clade when including these genomes is 1.7, far higher than species-level designations (Table S8). The final *de novo* assembled ancient genome from Germany falls in a clade with the *T. serpentiformis* genome. High bootstrap values on deep nodes in the tree (Figure 4B) support the separation of these *Tannerella* genomes into distinct clades.

The *Tannerella* genome from Germany that falls near *T. serpentiformis* is from the medieval G12 individual published by [12]. This study reported the presence of multiple *Tannerella* species, further resolution was not possible due to the genome availability of only one species, *T. forsythia*, within the *Tannerella* genus at the time. To investigate whether the assembled *Tannerella* MAG from this sample is *T. serpentiformis*, we further mapped all G12 sample libraries against the *T. serpentiformis* reference genome. We observed similar damage patterns when mapping to either the *T. forsythia* or the *T. serpentiformis* genome (Figure S3), with slightly more mismatches detected when reads are mapped against the *T. serpentiformis* genome. This suggests that *T. serpentiformis* is not necessarily a closer match to the *Tannerella* species found in sample G12, and it is possible that this *Tannerella* genome represents another novel, as-yet-undescribed species in the genus.

To assess the pathogenic potential of the reconstructed genomes in the broader *Tannerella* clade, we examined a selection of eight known virulence factors for *T. forsythia* (Figure 4C) using a previously curated list [14]. The *T. forsythia* genome from Richard III is the only reconstructed MAG to include both surface layer (S-layer) glycoprotein genes *tfsA* and *tfsB*. While these virulence factor genes are nearly ubiquitous in the reference genomes, several could not be detected in the *de novo* reconstructed genomes (Figure 4B), which may be due to the lower completeness of these genomes rather than a true absence. For genomes that fall outside the main *T. forsythia* clade, a third or more of the virulence factor genes could not be detected. While this also may be due to a lower completeness of the reconstructed MAGs, it may also reflect a true absence given that these genes were not consistently detected in reference genomes and may not be ubiquitous in *Tannerella* species other than *T. forsythia*. Overall, additional exploration of the genomic and genetic diversity of oral *Tannerella* is warranted to better understand the phylogenetic relationships and gene content observed in this study.

## Discussion

We report the oral microbiome profile of four dental calculus samples from King Richard III of England, as well as new comparative dental calculus data from Neolithic and medieval Germany. DNA preservation within the dental calculus of Richard III was exceptional, but the reconstructed oral microbiota did not stand out from dental calculus of similar age and geographic origin regarding the number of microbial species detected, nor the abundance and distribution of microbial species. Despite the high DNA recovery and deep sequencing of the dental calculus metagenomes, no authentic dietary DNA was identified. Phylogenetic assessment of the abundant periopathogen *Tannerella forsythia* revealed that the *T. forsythia* genome reconstructed from Richard III is highly similar to genomes found in present-day living populations, as well as in other archaeological dental calculus from Northern Europe.

A wide range of dietary microfossils, proteins, and metabolites have been previously reported in modern and archaeological dental calculus [13,47–49], but dietary DNA is less well studied. Nevertheless, detection and authentication of dietary DNA in ancient dental calculus remains a potential source of information about diet at individual level. While DNA sequences derived from particular meat and vegetable sources have been reported in ancient dental calculus [12,50], few studies have reported on the presence or abundance of potential dietary reads, while those that have investigated dietary reads have generally been unsuccessful in authenticating them [20,21,51]. In some cases, dietary DNA may be present, but the overwhelming abundance of microbial reads reduces the chances of dietary reads being incorporated into DNA libraries and sequenced. It is also possible that dietary DNA is mostly degraded by enzymes in saliva [52] or in the microbial biofilm [53,54], and very little is incorporated into dental calculus.

Despite the deep sequencing of the dental calculus from Richard III, totaling nearly 400 million DNA sequences from three teeth, no authentic dietary DNA sequences were identified. Poor DNA preservation is unlikely to be the reason given the very high DNA recovery, which was similar to that from present-day dental calculus, and the excellent preservation of the oral microbiome. Additional or deeper sequencing may eventually produce enough additional DNA sequences to identify potential dietary components, but this approach is expensive and not guaranteed to produce results. The low number of putative dietary sequences observed in present-day dental plaque suggests that the incorporation and preservation of dietary DNA may be limited in oral biofilms. It is unclear why other dietary remains, such as plant microfossils and dietary proteins, are more robustly recovered from dental calculus, but at present it appears that dental calculus may not be a sufficient source of dietary DNA even when exceptionally well-preserved and deeply sequenced.

Isotopic analysis of femur and rib fragments from the skeleton of king Richard III suggest that he consumed a diet rich in non-local water sources, such as wine, and high trophic level game animals, such as fish and waterfowl, during the last 2-3 years of his life [9]. However, this aristocratic and unusually rich diet does not appear to have resulted in detectable microbial community changes compared to the dental calculus of lesser elites and commoners from across northern Europe, as all dental calculus analyzed in this study exhibited shared and overlapping species diversity. Whether the king’s diet may have promoted the growth of specific individual oral microbial species, rather than affecting the entire microbial community, is unclear. Microbial databases remain incomplete and insufficient to fully resolve many oral taxa at the species level, as evidenced by our findings for the genus *Tanerella*. To date, a direct relationship between diet and dental plaque biofilm species composition remains weakly and inconsistently supported in the literature [22,28]. Population-level dental calculus data collected over time in a single region are needed to more finely track species and strain-level microbial changes following a dietary change. Further advances in laboratory techniques that enable DNA extraction from distinct layers of microbial accumulation on dental calculus may also open the field to exploring oral microbiome changes throughout the lifetime of an individual.

*Tannerella forsythia* is an oral bacterial species strongly associated with periodontal disease in living populations [55,56], and it is grouped together with *Porphyromonas gingivalis* and *Treponema denticola* within the “red complex” consortium of oral species with high pathogenic potential [38]. Our analysis of virulence genes within reconstructed *T. forsythia* genomes from ancient dental calculus found that these genes have been present in the species for at least several centuries, confirming earlier mapping-based assessments [17–19]. Although we did not detect all eight genes in each reconstructed genome, this absence could in part be an artefact of *de novo* metagenome assembly. Current approaches rely on sufficient read length and coverage depth for genome reconstruction, and ancient DNA libraries may fall short of these lower limits [57].

Oral species diversity within the genus *Tannerella* has not been extensively explored. At present, only two named *Tannerella* species, *T. forsythia* and *T. serpentiformis*, are available in NCBI databases, but two additional unnamed species are recognized in the extended Human Oral Microbiome Database (eHOMD) [44,45]. These two unnamed species, designated HMT-808 and HMT-916, do not have available genomes. As we downloaded only named *Tannerella* genomes from NCBI for our analysis, we do not expect our tree to include these additional species. However, our *Tannerella* phylogeny and virulence factor analysis hint that historic dental calculus samples contain *Tannerella* species other than *T. forsythia* and *T. serpentiformis*. In our phylogeny, 29 *Tannerella* genomes–including 3 historic Dutch genomes, 4 historic English genomes, and the genome from King Richard III–fall into a large clade that appears to represent true *T. forsythia* genomes. Only a single *Tannerella* genome from medieval Germany appears to belong to *T. serpentiformis* (or a closely related species). However, two historic English and 1 Neolithic German *Tannerella* genomes, as well as 2 NCBI genomes designated *T. forsythia*, fall basal to the main *T. forsythia* clade, suggesting that these genomes may represent a distinct *Tannerella* species that is more similar to *T. forsythia* than to *T. serpentiformis*. While patristic distance calculations support this finding, further genomic investigation is necessary to confirm these observations.

In addition to discovering additional *Tannerella* species diversity, our data suggest higher *Tannerella* diversity within the dental calculus of specific individuals. The elevated baseline damage frequency observed when mapping *Tannerella* DNA sequences from Richard III and other ancient individuals from England and Germany against the reference *T. forsythia* genome supports that these calculus samples may contain more than one *Tannerella* species. Because only a single *Tannerella* genome was reconstructed from each sample, however, it is likely that one species is dominant and present at an abundance that enabled *de novo* genome assembly, while the other species are less abundant and did not have sufficient sequences for *de novo* assembly [12,57]. While several studies have performed genomic and genetic investigations of *T. forsythia* in ancient dental calculus [12,17–19,58], they did not consider how the presence of other unnamed or unknown *Tannerella* species may have affected their analyses. Further investigation may reveal that some of the reported patterns of *T. forsythia* virulence gene prevalence in past populations can be attributed to these additional species.

## 5 Conclusion

The dental calculus of King Richard III is exceptionally well-preserved and ancient DNA analysis has revealed an oral microbiome species composition that is highly similar to that of other human dental calculus from northern Europe. Despite excellent DNA preservation and deep sequencing of the DNA libraries, we were unable to identify authentic dietary DNA. We identified a higher level of taxonomic diversity in *de novo* reconstructed *Tannerella* genomes than has been reported to date in ancient dental calculus, indicating that further investigation of the diversity of the periodontal disease-associated genus *Tannerella* is warranted to better understand how the oral microbiome has changed through human history.

## Methods

### Dental calculus sampling

Dental calculus samples of King Richard III were collected from lingual surfaces of three teeth: left lower first premolar, left lower canine, and right lower second premolar. Prior to analysis, the dental calculus from the canine was split into two sub-fractions, resulting in a total of 4 samples: R3-1 (GRF001.A), R3-2A (GRF001.B), R3-2B (GRF001.C), and R3-3 (GRF001.D) (Table 1; Table S1A). Six swabs from the archaeological storage facility were collected to monitor for possible contamination: sample bag (R3-Bag), sample box (R3-Box), storage refrigerator (R3-Fr), laboratory gloves (R3-Gl), osteologist hands (R3-HS), and a laboratory water sample (R3MODBL) (Table S1A). Samples and swabs were transferred to the ancient DNA cleanroom facilities at the Laboratories of Molecular Anthropology and Microbiome Research (LMAMR) at the University of Oklahoma for DNA extraction.

Dental calculus of three additional individuals from the Neolithic Linearbandkeramik (LBK, 6th millennium BCE) site of Stuttgart-Mühlhausen (Viesenhäuser Hof) in Baden-Württemberg, Germany [32,59–61] were also collected for comparative analysis (Table S1A): gr. I-064/335 (SMH005.A), gr. I-056/528 (SMH007.A), gr. II-027/274 (SMH008.A). Samples were transferred to the ancient DNA cleanroom facilities at the University of Tübingen, Germany for DNA extraction.

### DNA extraction

DNA from the dental calculus of Richard III was extracted in a dedicated ancient DNA facility at LMAMR in accordance with established contamination control precautions and workflows. Two cleanroom non-template extraction controls (R3-AncBL1, R3-AncBL2) were processed alongside the experimental samples during all analytical steps to monitor for possible contamination. Prior to decalcification, dental calculus samples were UV-irradiated for 2 min using a Stratagene Stratalinker UV 1800 Crosslinker. To remove remaining surface debris and contaminants, they were agitated in 1 ml of 0.5 M EDTA solution for 15 min and decanted. The dental calculus samples were then processed using the DNA extraction protocol described in [62], with minor modifications. In brief, the samples were crushed to powder using a sterile steel spatula and resuspended in 1 ml 0.5M EDTA (Sigma) and incubated overnight at room temperature. A 100 μl proteinase K solution (>600 mAU ml^-1^; Qiagen) was then added and incubated at 37°C for 8 hours, followed by continued digestion under agitation at room temperature until decalcification was complete. The DNA was then purified using a MinElute column (Qiagen) connected to a Zymo reservoir (Zymo Research) following modified manufacturer instructions in which the amount of PB binding buffer was increased to 13 ml [62]. The DNA was eluted in 60 μl EB buffer (Qiagen), and the concentration of the eluate was quantified using 1 μl with a Qubit HS assay (Life Technologies).

DNA from the Stuttgart-Mühlhausen dental calculus samples was extracted in a dedicated ancient DNA facility at the University of Tübingen in accordance with established contamination control precautions and workflows, and following the DNA extraction protocol described in [62] without modification.

### Illumina library construction

Shotgun Illumina libraries (Table S1B) were constructed from 60-100 ng DNA from Richard III calculus samples and 30 μl of extract for non-template extraction controls using a NEBNext DNA Library Prep Master set for 454 (New England Biolabs, E6070) according to manufacturer instructions. End repair was performed in 50 μl reactions with 100 ng of DNA. The end repair cocktail was incubated for 20 min at 12°C and 15 min at 37°C and then purified using a MinElute column (Qiagen) and eluted in 30 μl EB buffer. Blunt end adapters (IS1/IS3 and IS2/IS3) were prepared following Meyer and Kircher (2010) and ligated to the end-repaired DNA in 50μl reactions. The reaction was incubated for 15 min at 20°C and then purified using QiaQuick columns (Qiagen) and eluted in 30 μl EB. The adapter fill-in reaction was performed in a final volume of 50 μl and incubated for 20 min at 37°C followed by 20 min at 80°C to inactivate the Bst polymerase. Libraries were amplified and indexed in a 50 μl PCR reaction, using 26.3 μl H_2_O, 5 μl 10x Platinum Taq MasterMix, 3 μl 2mM dNTPs, 1 μl BSA (2.5 mg/ml), 1.5 μl 50 mM MgCl2, 1.5 μl i5 indexing primer (10 μM), 1.5 μl i7 indexing primer (10 μM), 0.2 μl PlatinumTaq, and 10μl of library template. Thermocycling conditions were 2 min at 95°C, followed by 12 cycles of 15 s at 95°C, 30 s at 60°C, and 30 s at 72°C, and a final 7 min elongation step at 72°C. The optimal number of PCR cycles for each sample was first confirmed by qPCR, and all negative controls were amplified for 20 cycles. All libraries were purified using a MinElute PCR Purification kit (Qiagen) following manufacturer instructions and quantified using a Bioanalyzer 2100 High Sensitivity DNA assay (Agilent).

Shotgun Illumina libraries (Table S1B) were prepared similarly for the Stuttgart-Mühlhausen dental calculus samples, with minor modifications as described in the Non-UDG Treated Double-Stranded Ancient DNA Library Preparation for Illumina Sequencing protocol available on protocols.io (dx.doi.org/10.17504/protocols.io.bakricv6). A second aliquot of extracted DNA was then additionally prepared following USER enzyme treatment in order to remove DNA damage, as described in the Full-UDG Treated Double-Stranded Ancient DNA Library Preparation for Illumina Sequencing protocol available on protocols.io (dx.doi.org/10.17504/protocols.io.bqbpmsmn).

### DNA sequencing

Richard III DNA libraries (Table S1B) were pooled in equimolar amounts and sequenced on an Illumina HiSeq 2000 instrument with PE 2x100bp chemistry in RapidRun mode at the Yale Center for Genome Analysis (YCGA). A total of 393,365,842 sequencing reads were generated: R3-1, 73,678,628; R3-2A, 104,804,438; R3-2B, 125,374,688; R3-3, 89,508,088.

At the MPI-EVA, the Stuttgart-Mühlhausen non-UDG treated libraries (SMH005.A0101, SMH007.A0101, SMH008.A0101; Table S1B) were sequenced on an Illumina NextSeq 500 with PE 2x75bp chemistry, and the full-UDG libraries (SMH005.A0102, SMH007.A0102, SMH008.A0102; Table S1B) were sequenced on an Illumina HiSeq 4000 with PE 2x75bp chemistry. A total of 724,359,015 sequencing reads were generated: SMH005.A, 213,889,522; SMH007.A, 164,169,600; SMH008.A, 346,299,893.

To generate more medieval dental calculus sequencing data for comparative analysis, we also deeply sequenced four previously generated DNA libraries (Table S1B) of individual G12 (S5, S6, S7, S8) from the site of Dalheim, Germany [12] on two flow cells of an Illumina HiSeq 2000 using PE 2x150bp chemistry at the Yale Center for Genome Analysis (YCGA). A total of 240,767,032 sequencing reads were generated: S5, 86,797,120; S6, 55,998,288; S7, 50,206,795; S8, 47,764,829.

### Data processing

Raw data was processed using the nf-core/eager v2.4.4 pipeline [63], including adapter trimming, read quality-filtering (min length 30bp, quality 40), and merging with adapter-removal, mapping against human genome hg19 with bwa aln v0.7.17-r1188 [64], and selecting all reads that did not map to the human genome into a single fasta file with samtools v1.17 [65]. These fasta files of non-human reads were used for taxonomic profiling.

### Published comparative samples

Normalized DNA extraction yields (ng DNA/mg calculus) measured in our laboratory from 143 archaeological dental calculus samples from diverse global sites and 3 present-day individuals (Figure 1A) were obtained from the literature [11,12,29,31,51,66–68] and are provided in Table S2.

Published dental calculus metagenomic datasets from northern Europe dating 200-1000 BP were selected using AncientMetagenomeDir (accessed 2024/01) for [69] (Table S3). These included samples from England [30,70,71], Ireland [29], the Netherlands [31], and Germany [12]. All sequencing data for these samples were downloaded from ENA and processed with the nf-core/eager pipeline as described above for the new genetic data generated in this study, but datasets with fewer than 1M sequenced reads were excluded due to high stochasticity in identified species obfuscating signals of contamination or false-positive identification vs. true presence. This approach excluded all environmental swabs (R3-AncBL1, R3-AncBL2, R3Bag, R3Box, R3FR, R3GL, R3HS, R3MODBL) collected during sampling of the four calculus samples from Richard III.

### Taxonomic profiling and diversity analysis

Newly sequenced and published metagenomic datasets ranged from <3M to >300M reads (Figure 1B), with most datasets between 5M and 30M reads. Notable exceptions were the newly sequenced libraries from King Richard III and from Germany (Stuttgart-Mühlhausen and Dalheim), which were intentionally deeply sequenced to facilitate *de novo* genome assembly. To improve data quality and consistency, samples with fewer than 5M reads were excluded from all downstream analysis. All samples were taxonomically profiled with kraken2 v2.1.3 [33,34] using the GTDB v202 [35] database downloaded from the Struo2 [72] ftp server (http://ftp.tue.mpg.de/ebio/projects/struo2/). Taxonomic tables were filtered in R v4.3.2 to remove potential contamination by removing all species with fewer than 5,000 reads across all samples and then filtering out all species that were present at less than 0.01% relative abundance. The number of species and Shannon index were calculated in R using the package vegan v2.6-8 [73]. A principal component analysis (PCA) was run in R with the package mixOmics v6.26.0 [74]. After adding a value of +1 to all entries in the table to remove 0 values, a centered-log ratio transformation [75] was run on the dataset and a PCA was performed on the transformed datatable. All diversity analysis results were plotted in R with ggplot2 [76].

### Microbiome preservation assessment

Preservation of the samples was assessed using two methods. First, SourceTracker v1.0 [77] was applied to the newly generated data to determine whether the species profile was predominantly attributable to an oral source or to potential contamination sources (Table S4). Comparative sources included modern human dental calculus [14,30], dental plaque [78], stool [78–80], and skin [78,81], as well as archaeological bone [14] and sediment [82]. Second, the R package cuperdec [14] was used to assess the proportion of taxa from an oral source within all samples, including both newly generated sequences and published comparative data. Negative controls and one published dental calculus sample failed to meet the threshold for high oral content (Table S2; Table S3) and were excluded.

### Dietary analysis

For identification of putative dietary DNA sequences, we used the high-throughput aligner MALT [36,37] together with the NCBI nt database (October 2017; uploaded to Zenodo under DOI: 10.5281/zenodo.4382154) to taxonomically profile: the Richard III dental calculus samples (R3-1, R3-2A, R3-2B, and R3-3), negative controls (R3-AncBL1, R3-AncBL2), and environmental controls (R3Bag, R3Box, R3FR, R3GL, R3HS, R3MODBL); and the dental plaque [78], stool [78–80], skin [78,81], archaeological bone [14] and archaeological sediment [82] datasets also used for preservation assessment. We employed a relaxed percent identity parameter of 85% and a base tail cut off (“minimum support”) of 0.01%. Resulting RMA6 files were loaded into MEGAN6 CE [83] and Operational Taxonomic Unit (OTU) tables were exported for Spermatophyta (seed plants) and Euteleostomi (major clade of vertebrates) (Table S5).

### De novo genome reconstruction

All metagenome samples were *de novo* assembled and binned using a Snakemake [84] pipeline previously described in [85], and which is available on GitHub under https://github.com/alexhbnr/ancient_metagenome_assembly. Assemblies were performed on an individual level, such that datasets from all libraries of a single individual were assembled together, resulting in a single output of metagenome-assembled genomes from all four libraries of Richard III. In brief, unmerged paired-end sequencing datasets were assembled using MEGAHIT v1.2.9 [86]. After correcting the consensus contig sequences to remove miscoding lesions due to the presence of ancient DNA [85], the contigs were binned using metaBAT v2.15 [87], MaxBin v2.2.7 [88], and CONCOCT v1.1.0 [89] and subsequently refined using metaWRAP v1.3.2 [90]. The resulting metagenome-assembled genomes (MAGs) (Table S6) were further refined and validated and annotated with bakta v1.9.4 [91] using a Snakemake pipeline previously described in [85] and available on GitHub under https://github.com/alexhbnr/automatic_MAG_refinement to remove potentially chimeric contigs.

### Phylogenetic analysis and authentication

All genomes designated as *Tannerella forsythia* in NCBI databases were downloaded for phylogenetic comparison to the *de novo* reconstructed ancient genomes. The completeness and contamination of all reference genomes from NCBI were assessed with checkM v1.2.2 [92] using the program dRep v3.4.3 [93] (Table S6). A genome of *Tannerella serpentiformis* (GCA_003033925.1) was downloaded for use as an outgroup to root the tree. All NCBI genomes and all *de novo* reconstructed ancient genomes were used as input for the program PhyloPhlAn v3.1 [94,95] to reconstruct a tree based on selected marker genes specific for the species *T. forsythia*. Bootstrapping (200 replicates) was performed by RAxML using RAxML-NG v1.2.2 [96] on the protein marker gene alignment file produced by Phylophlan. The resulting tree was loaded into R and visualized with ggtree [97] with a midpoint root. Patristic distances were calculated from the tree using cophenetic.phylo in the R package ape v5.8 [98] (Table S8).

The reconstructed genomes were confirmed to be of ancient origin by assessing the damage patterns on reads from each sample mapped against the *T. forsythia* reference genome (NC_016610.1). All libraries were mapped using bwa aln v0.7.17-r1188 with the flags -n 0.01 -o 2 -l 16500. Mapped reads were filtered with samtools v1.17, and only reads with a mapping quality of ≥ 25 and a length ≥ 30bp were considered suitable for damage analysis (Table S7). DamageProfiler v1.1 [99] was used to quantify DNA damage on mapped reads. Libraries of the sample G12 were additionally mapped against the *T. serpentiformis* reference genome, and the mapped reads were also filtered and analyzed using DamageProfiler.

### Virulence gene content assessment

The *Tanerella* genomes downloaded from NCBI were annotated with bakta [91] using the same settings as in the *de novo* assembly pipeline above. The annotated gene names were cross-referenced against the known *T. forsythia* virulence factors listed in [14], and marked as 1 if present and 0 if absent (Table S6). The binary table was loaded into R and visualized as a matrix using tidyverse v1.3.0 [100] and ggplot2 [76], and then combined with the phylogenetic tree using patchwork v1.2.0 [101]. The completeness and contamination of all genomes were likewise visualized in the same way using R.

### Dietary DNA assessment

Potential dietary DNA sequences were assessed using MALT [36,37]. The metagenomic datasets were aligned with MALT v 0.4.0 against the NCBI nt database (October 2016). Reads matching eukaryotes were evaluated for the number of reads mapped and damage patterns to identify genuine ancient reads. No potential dietary eukaryotes could be confirmed as authentic due to insufficient numbers of reads or a lack of DNA damage patterns.

## Author Contributions

T.K., A.R., and C.W. conceived the study. T.K., E.R., C.M.L., M.F., J.W., J.K., and C.W. provided samples and resources. A.M., C.H., C.S., S.F., and A.O. performed lab work. I.M.V., A.H., Z.F., and J.F.Y., performed data analysis and interpretation. I.V. and C.W. wrote the manuscript with input from all authors.

## Acknowledgments

We thank Joanna Drath, Eva Rosenstock, Alisa Hujic, and Michal Feldman for assistance with the Stuttgart-Mühlhausen remains. Funding for this study was provided by the Werner Siemens Stiftung (“Paleobiotechnology” to C.W.), the Deutsche Forschungsgemeinschaft (DFG, German Research Foundation) under Germany’s Excellence Strategy (EXC 2051 Project-ID 390713860, “Balance of the Microverse”), the Max Planck Harvard Research Center for the Archaeoscience of the Ancient Mediterranean (MHAAM), and the Max Planck Society.

## Ethics statement

All required permissions were obtained to perform genetic data generation from newly analyzed archaeological dental calculus in this study. The excavation and analysis of the remains of Richard III was carried out by the University of Leicester Archaeological Services as part of the excavation of the Grey Friars Friary under a licence granted by the U.K Ministry of Justice. Permission to analyze remains from the site of Stuttgart-Mühlhausen was provided by the State Office for Monument Preservation in Konstanz, Germany.

## Data Availability Statement

All new sequencing data generated for this study was deposited in the European Nucleotide Archive under accession PRJEB84038 at https://www.ebi.ac.uk/ena/browser/view/PRJEB84038. Code used to perform analyses and generate the figures is provided in a GitHub repository at https://github.com/ivelsko/RIII_oral_micro.

## References

1. Buckley R, Morris M, Appleby J, King T, O’Sullivan D, Foxhall L. 2013 ‘The king in the car park’: new light on the death and burial of Richard III in the Grey Friars church, Leicester, in 1485. Antiquity 87, 519–538.

2. Appleby J et al. 2015 Perimortem trauma in King Richard III: a skeletal analysis. Lancet 385, 253–259.

3. Appleby J, Mitchell PD, Robinson C, Brough A, Rutty G, Harris RA, Thompson D, Morgan B. 2014 The scoliosis of Richard III, last Plantagenet King of England: diagnosis and clinical significance. Lancet 383, 1944.

4. King TE et al. 2014 Identification of the remains of King Richard III. Nat. Commun. 5, 5631.

5. Rous J. 1716 Historia regum Angliae. Oxonii.

6. More T. 1883 More’s History of King Richard III. Cambridge, England: Cambridge University Press.

7. Shakespeare W. 1800 Richard III, a tragedy. Manchester: R. & W. Dean & Co.

8. Baldwin D. 2015 Richard III. Oxford, England: Amberley Publishing.

9. Lamb AL, Evans JE, Buckley R, Appleby J. 2014 Multi-isotope analysis demonstrates significant lifestyle changes in King Richard III. J. Archaeol. Sci. 50, 559–565.

10. Velsko IM, Warinner CG. 2017 Bioarchaeology of the human microbiome. Bioarchaeology International 1, 86–99.

11. Warinner C, Speller C, Collins MJ. 2015 A new era in palaeomicrobiology: prospects for ancient dental calculus as a long-term record of the human oral microbiome. Philos. Trans. R. Soc. Lond. B Biol. Sci. 370, 20130376.

12. Warinner C et al. 2014 Pathogens and host immunity in the ancient human oral cavity. Nat. Genet. 46, 336–344.

13. Scott A et al. 2021 Exotic foods reveal contact between South Asia and the Near East during the second millennium BCE. Proc. Natl. Acad. Sci. U. S. A. 118, e2014956117.

14. Fellows Yates JA et al. 2021 The evolution and changing ecology of the African hominid oral microbiome. Proc. Natl. Acad. Sci. U. S. A. 118. (doi:10.1073/pnas.2021655118)

15. Velsko IM, Warinner C. 2025 Streptococcus abundance and oral site tropism in humans and non-human primates reflects host and lifestyle differences. NPJ Biofilms Microbiomes 11, 19.

16. Granehäll L et al. 2021 Metagenomic analysis of ancient dental calculus reveals unexplored diversity of oral archaeal Methanobrevibacter. Microbiome 9, 197.

17. Bravo-Lopez M et al. 2020 Paleogenomic insights into the red complex bacteria Tannerella forsythia in Pre-Hispanic and Colonial individuals from Mexico. Philos. Trans. R. Soc. Lond. B Biol. Sci. 375, 20190580.

18. Philips A et al. 2020 Analysis of oral microbiome from fossil human remains revealed the significant differences in virulence factors of modern and ancient Tannerella forsythia. BMC Genomics 21, 402.

19. Jackson I, Woodman P, Dowd M, Fibiger L, Cassidy LM. 2024 Ancient genomes from bronze age remains reveal deep diversity and recent adaptive episodes for human oral pathobionts. Mol. Biol. Evol. 41. (doi:10.1093/molbev/msae017)

20. Gancz AS et al. 2023 Ancient dental calculus reveals oral microbiome shifts associated with lifestyle and disease in Great Britain. Nat. Microbiol. 8, 2315–2325.

21. Mann AE, Fellows Yates JA, Fagernäs Z, Austin RM, Nelson EA, Hofman CA. 2023 Do I have something in my teeth? The trouble with genetic analyses of diet from archaeological dental calculus. Quat. Int. 653-654, 33–46.

22. Velsko IM, Gallois S, Stahl R, Henry AG, Warinner C. 2023 High conservation of the dental plaque microbiome across populations with differing subsistence strategies and levels of market integration. Mol. Ecol. (doi:10.1111/mec.16988)

23. Kato I, Vasquez A, Moyerbrailean G, Land S, Djuric Z, Sun J, Lin H-S, Ram JL. 2017 Nutritional correlates of human oral microbiome. J. Am. Coll. Nutr. 36, 88–98.

24. Lokmer A et al. 2020 Response of the human gut and saliva microbiome to urbanization in Cameroon. Sci. Rep. 10, 2856.

25. Lassalle F, Spagnoletti M, Fumagalli M, Shaw L, Dyble M, Walker C, Thomas MG, Bamberg Migliano A, Balloux F. 2018 Oral microbiomes from hunter-gatherers and traditional farmers reveal shifts in commensal balance and pathogen load linked to diet. Mol. Ecol. 27, 182–195.

26. Ottoni C et al. 2021 Tracking the transition to agriculture in Southern Europe through ancient DNA analysis of dental calculus. Proc. Natl. Acad. Sci. U. S. A. 118. (doi:10.1073/pnas.2102116118)

27. Quagliariello A et al. 2022 Ancient oral microbiomes support gradual Neolithic dietary shifts towards agriculture. Nat. Commun. 13, 6927.

28. Innocenti G, Martino ME, Stellini E, Di Fiore A, Quagliariello A. 2023 Dental calculus microbiome correlates with dietary intake. Mol. Oral Microbiol. 38, 189–197.

29. Mann AE et al. 2018 Differential preservation of endogenous human and microbial DNA in dental calculus and dentin. Sci. Rep. 8, 9822.

30. Velsko IM et al. 2019 Microbial differences between dental plaque and historic dental calculus are related to oral biofilm maturation stage. Microbiome 7, 102.

31. Velsko IM et al. 2022 Ancient dental calculus preserves signatures of biofilm succession and interindividual variation independent of dental pathology. PNAS Nexus 1, gac148.

32. Price TD, Wahl J, Knipper C, Burger-Heinrich E, Kurtz G, Ra} B. 2003 Das bandkeramische Gräberfeld vom „Viesenhäuser Hof “bei Stuttgart-Mühlhausen: Neue Untersuchungsergebnisse zum Migrationsverhalten im frühen Neolithikum. Kommissionsverlag Konrad Theiss Verlag

33. Wood DE, Lu J, Langmead B. 2019 Improved metagenomic analysis with Kraken 2. Genome Biol. 20, 257.

34. Lu J, Rincon N, Wood DE, Breitwieser FP, Pockrandt C, Langmead B, Salzberg SL, Steinegger M. 2022 Metagenome analysis using the Kraken software suite. Nat. Protoc. 17, 2815–2839.

35. Parks DH, Chuvochina M, Rinke C, Mussig AJ, Chaumeil P-A, Hugenholtz P. 2022 GTDB: an ongoing census of bacterial and archaeal diversity through a phylogenetically consistent, rank normalized and complete genome-based taxonomy. Nucleic Acids Res. 50, D785– D794.

36. Vågene ÅJ et al. 2018 Salmonella enterica genomes from victims of a major sixteenth-century epidemic in Mexico. Nat Ecol Evol 2, 520–528.

37. Herbig A, Maixner F, Bos KI, Zink A, Krause J, Huson DH. 2016 MALT: Fast alignment and analysis of metagenomic DNA sequence data applied to the Tyrolean Iceman. bioRxiv., 050559. (doi:10.1101/050559)

38. Socransky SS, Haffajee AD, Cugini MA, Smith C, Kent RL Jr. 1998 Microbial complexes in subgingival plaque. J. Clin. Periodontol. 25, 134–144.

39. Bowers RM et al. 2017 Minimum information about a single amplified genome (MISAG) and a metagenome-assembled genome (MIMAG) of bacteria and archaea. Nat. Biotechnol. 35, 725–731.

40. Briggs AW et al. 2007 Patterns of damage in genomic DNA sequences from a Neandertal. Proc. Natl. Acad. Sci. U. S. A. 104, 14616–14621.

41. Rohland N, Harney E, Mallick S, Nordenfelt S, Reich D. 2015 Partial uracil–DNA– glycosylase treatment for screening of ancient DNA. Philos. Trans. R. Soc. Lond. B Biol. Sci. 370, 20130624.

42. Briggs AW, Stenzel U, Meyer M, Krause J, Kircher M, Pääbo S. 2010 Removal of deaminated cytosines and detection of in vivo methylation in ancient DNA. Nucleic Acids Res. 38, e87.

43. Warinner C, Herbig A, Mann A, Fellows Yates JA, Weiß CL, Burbano HA, Orlando L, Krause J. 2017 A Robust Framework for Microbial Archaeology. Annu. Rev. Genomics Hum. Genet. 18, 321–356.

44. Chen T, Yu W-H, Izard J, Baranova OV, Lakshmanan A, Dewhirst FE. 2010 The Human Oral Microbiome Database: a web accessible resource for investigating oral microbe taxonomic and genomic information. Database (Oxford) 2010, baq013.

45. Escapa IF, Chen T, Huang Y, Gajare P, Dewhirst FE, Lemon KP. 2018 New insights into human nostril microbiome from the expanded Human Oral Microbiome Database (eHOMD): A resource for the microbiome of the human aerodigestive tract. mSystems 3. (doi:10.1128/mSystems.00187-18)

46. Velsko IM, Perez MS, Richards VP. 2019 Resolving Phylogenetic Relationships for Streptococcus mitis and Streptococcus oralis through Core- and Pan-Genome Analyses. Genome Biol. Evol. 11, 1077–1087.

47. Velsko IM et al. 2017 The dental calculus metabolome in modern and historic samples. Metabolomics 13, 134.

48. Hendy J et al. 2018 Proteomic evidence of dietary sources in ancient dental calculus. Proc. Biol. Sci. 285, 20180977.

49. Radini A, Nikita E. 2023 Beyond dirty teeth: Integrating dental calculus studies with osteoarchaeological parameters. Quat. Int. 653-654, 3–18.

50. Weyrich LS et al. 2017 Neanderthal behaviour, diet, and disease inferred from ancient DNA in dental calculus. Nature. 544, 357–361. (doi:10.1038/nature21674)

51. Velsko IM et al. 2024 Exploring the potential of dental calculus to shed light on past human migrations in Oceania. Nat. Commun. 15, 10191.

52. Bradbury S. 1956 Human saliva as a convenient source of ribonuclease. J. Cell Sci. S3-97, 323–327.

53. Liu J, Sun L, Liu W, Guo L, Liu Z, Wei X, Ling J. 2017 A nuclease from Streptococcus mutans facilitates biofilm dispersal and escape from killing by neutrophil extracellular traps. Front. Cell. Infect. Microbiol. 7, 97.

54. Rostami N et al. 2022 Interspecies competition in oral biofilms mediated by Streptococcus gordonii extracellular deoxyribonuclease SsnA. NPJ Biofilms Microbiomes 8, 96.

55. Schäffer C, Andrukhov O. 2024 The intriguing strategies of Tannerella forsythia’s host interaction. Front. Oral Health 5, 1434217.

56. Ready D, D’Aiuto F, Spratt DA, Suvan J, Tonetti MS, Wilson M. 2008 Disease severity associated with presence in subgingival plaque of Porphyromonas gingivalis, Aggregatibacter actinomycetemcomitans, and Tannerella forsythia, singly or in combination, as detected by nested multiplex PCR. J. Clin. Microbiol. 46, 3380–3383.

57. Borry M, Hübner A, Rohrlach AB, Warinner C. 2021 PyDamage: automated ancient damage identification and estimation for contigs in ancient DNA de novo assembly. PeerJ 9, e11845.

58. Honap TP et al. 2023 Oral metagenomes from Native American Ancestors reveal distinct microbial lineages in the pre-contact era. Am J Biol Anthropol (doi:10.1002/ajpa.24735)

59. Cox SL, Nicklisch N, Francken M, Wahl J, Meller H, Haak W, Alt KW, Rosenstock E, Mathieson I. 2024 Socio-cultural practices may have affected sex differences in stature in Early Neolithic Europe. Nat. Hum. Behav. 8, 243–255.

60. Ash A, Francken M, Pap I, Tvrdý Z, Wahl J, Pinhasi R. 2016 Regional differences in health, diet and weaning patterns amongst the first Neolithic farmers of central Europe. Sci. Rep. 6, 29458.

61. Rivollat M et al. 2020 Ancient genome-wide DNA from France highlights the complexity of interactions between Mesolithic hunter-gatherers and Neolithic farmers. Sci. Adv. 6, eaaz5344.

62. Dabney J et al. 2013 Complete mitochondrial genome sequence of a Middle Pleistocene cave bear reconstructed from ultrashort DNA fragments. Proc. Natl. Acad. Sci. U. S. A. 110, 15758–15763.

63. Fellows Yates JA, Lamnidis TC, Borry M, Andrades Valtueña A, Fagernäs Z, Clayton S, Garcia MU, Neukamm J, Peltzer A. 2021 Reproducible, portable, and efficient ancient genome reconstruction with nf-core/eager. PeerJ 9, e10947.

64. Li H, Durbin R. 2009 Fast and accurate short read alignment with Burrows-Wheeler transform. Bioinformatics 25, 1754–1760.

65. Danecek P et al. 2021 Twelve years of SAMtools and BCFtools. Gigascience 10. (doi:10.1093/gigascience/giab008)

66. Ziesemer KA et al. 2015 Intrinsic challenges in ancient microbiome reconstruction using 16S rRNA gene amplification. Sci. Rep. 5, 16498.

67. Ozga AT, Nieves-Colón MA, Honap TP, Sankaranarayanan K, Hofman CA, Milner GR, Lewis CM Jr, Stone AC, Warinner C. 2016 Successful enrichment and recovery of whole mitochondrial genomes from ancient human dental calculus. Am. J. Phys. Anthropol. 160, 220–228.

68. Standeven FJ et al. 2024 An extensive archaeological dental calculus dataset spanning 5000 years for ancient human oral microbiome research. bioRxiv. (doi:10.1101/2024.09.17.613443)

69. Fellows Yates JA et al. 2021 Community-curated and standardised metadata of published ancient metagenomic samples with AncientMetagenomeDir. Sci Data 8, 31.

70. Farrer AG, Wright SL, Skelly E, Eisenhofer R, Dobney K, Weyrich LS. 2021 Effectiveness of decontamination protocols when analyzing ancient DNA preserved in dental calculus. Sci. Rep. 11, 7456.

71. Fagernäs Z, García-Collado MI, Hendy J, Hofman CA, Speller C, Velsko I, Warinner C. 2020 A unified protocol for simultaneous extraction of DNA and proteins from archaeological dental calculus. J. Archaeol. Sci. 118, 105135.

72. Youngblut ND, Ley RE. 2021 Struo2: efficient metagenome profiling database construction for ever-expanding microbial genome datasets. PeerJ 9, e12198.

73. Oksanen, J. 2007 Vegan : community ecology package version 1.8-6. http://cran.r-project.org

74. Rohart F, Gautier B, Singh A, Lê Cao K-A. 2017 mixOmics: An R package for ‘omics feature selection and multiple data integration. PLoS Comput. Biol. 13, e1005752.

75. Gloor GB, Macklaim JM, Pawlowsky-Glahn V, Egozcue JJ. 2017 Microbiome datasets are compositional: And this is not optional. Front. Microbiol. 8. (doi:10.3389/fmicb.2017.02224)

76. Wickham H. 2016 ggplot2: Elegant Graphics for Data Analysis. Springer.

77. Knights D, Kuczynski J, Charlson ES, Zaneveld J, Mozer MC, Collman RG, Bushman FD, Knight R, Kelley ST. 2011 Bayesian community-wide culture-independent microbial source tracking. Nat. Methods 8, 761–763.

78. Human Microbiome Project Consortium. 2012 Structure, function and diversity of the healthy human microbiome. Nature 486, 207–214.

79. Obregon-Tito AJ et al. 2015 Subsistence strategies in traditional societies distinguish gut microbiomes. Nat. Commun. 6, 6505.

80. Rampelli S et al. 2015 Metagenome Sequencing of the Hadza Hunter-Gatherer Gut Microbiota. Curr. Biol. 25, 1682–1693.

81. Oh J, Byrd AL, Park M, NISC Comparative Sequencing Program, Kong HH, Segre JA. 2016 Temporal stability of the human skin microbiome. Cell 165, 854–866.

82. Slon V et al. 2017 Neandertal and Denisovan DNA from Pleistocene sediments. Science 356, 605–608.

83. Huson DH, Beier S, Flade I, Górska A, El-Hadidi M, Mitra S, Ruscheweyh H-J, Tappu R. 2016 MEGAN Community Edition - interactive exploration and analysis of large-scale microbiome sequencing data. PLoS Comput. Biol. 12, e1004957.

84. Mölder F et al. 2021 Sustainable data analysis with Snakemake. F1000Res. 10, 33.

85. Klapper M et al. 2023 Natural products from reconstructed bacterial genomes of the Middle and Upper Paleolithic. Science, eadf5300.

86. Li D, Liu C-M, Luo R, Sadakane K, Lam T-W. 2015 MEGAHIT: an ultra-fast single-node solution for large and complex metagenomics assembly via succinct de Bruijn graph. Bioinformatics 31, 1674–1676.

87. Kang DD, Li F, Kirton E, Thomas A, Egan R, An H, Wang Z. 2019 MetaBAT 2: an adaptive binning algorithm for robust and efficient genome reconstruction from metagenome assemblies. PeerJ 7, e7359.

88. Wu Y-W, Simmons BA, Singer SW. 2016 MaxBin 2.0: an automated binning algorithm to recover genomes from multiple metagenomic datasets. Bioinformatics 32, 605–607.

89. Alneberg J et al. 2014 Binning metagenomic contigs by coverage and composition. Nat. Methods 11, 1144–1146.

90. Uritskiy GV, DiRuggiero J, Taylor J. 2018 MetaWRAP-a flexible pipeline for genome-resolved metagenomic data analysis. Microbiome 6, 158.

91. Schwengers O, Jelonek L, Dieckmann MA, Beyvers S, Blom J, Goesmann A. 2021 Bakta: rapid and standardized annotation of bacterial genomes via alignment-free sequence identification. Microb Genom 7. (doi:10.1099/mgen.0.000685)

92. Parks DH, Imelfort M, Skennerton CT, Hugenholtz P, Tyson GW. 2015 CheckM: assessing the quality of microbial genomes recovered from isolates, single cells, and metagenomes. Genome Res. 25, 1043–1055.

93. Olm MR, Brown CT, Brooks B, Banfield JF. 2017 dRep: a tool for fast and accurate genomic comparisons that enables improved genome recovery from metagenomes through de-replication. ISME J. 11, 2864–2868.

94. Segata N, Börnigen D, Morgan XC, Huttenhower C. 2013 PhyloPhlAn is a new method for improved phylogenetic and taxonomic placement of microbes. Nat. Commun. 4, 2304.

95. Asnicar F et al. 2020 Precise phylogenetic analysis of microbial isolates and genomes from metagenomes using PhyloPhlAn 3.0. Nat. Commun. 11, 2500.

96. Kozlov AM, Darriba D, Flouri T, Morel B, Stamatakis A. 2019 RAxML-NG: a fast, scalable and user-friendly tool for maximum likelihood phylogenetic inference. Bioinformatics 35, 4453–4455.

97. Yu G, Smith DK, Zhu H, Guan Y, Lam TT-Y. 2017 Ggtree: An r package for visualization and annotation of phylogenetic trees with their covariates and other associated data. Methods Ecol. Evol. 8, 28–36.

98. Paradis E, Strimmer K, Claude J, Jobb G, Opgen-Rhein R, Dutheil J, Noel Y, Bolker B, Lemon J. 2008 The ape package. Analyses of phylogenetics and evolution

99. Neukamm J, Peltzer A, Nieselt K. 2021 DamageProfiler: fast damage pattern calculation for ancient DNA. Bioinformatics 37, 3652–3653.

100. Wickham H et al. 2019 Welcome to the Tidyverse. Journal of Open Source Software 4, 1686.

101. Pedersen TL. 2017 patchwork: The Composer of ggplots. R package version 0. 0 1.

