## Supplemental Figures S1-S3 for "The oral microbiome of King Richard III of England"

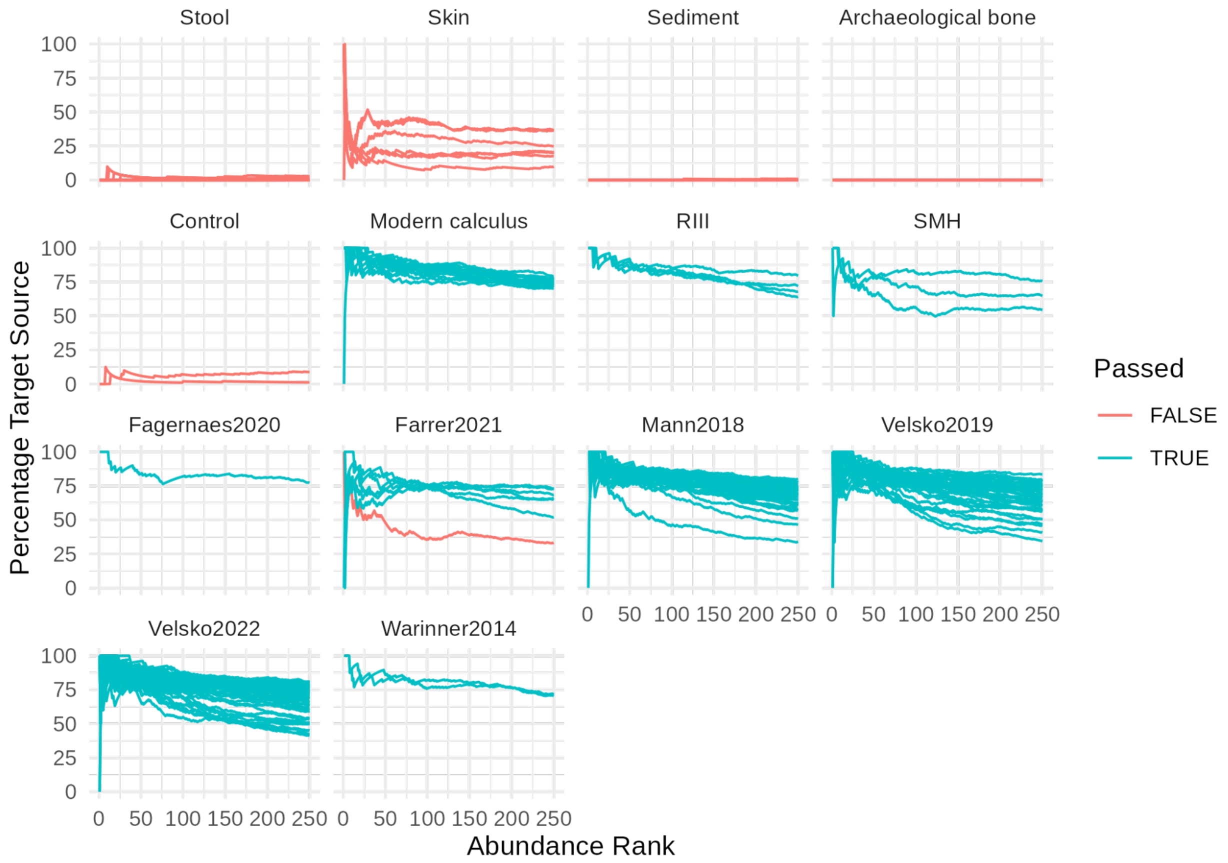


**Figure S1.** Cumulative percent decay (cuperdec) curves for ancient dental calculus samples including newly generated and published data as well as environmental controls (same as those used in SourceTracker) and extraction and library build blanks ('Control'). Samples are grouped by their original publication, titled by the first author's last name and the year of publication. Lines indicate the percent of all species (y-axis) at and above a given abundance rank (x-axis) that are classified as oral. Passed - False indicates the sample did not pass the preservation cut-off, as defined by the modern dental calculus and other control samples. Passed - True indicates the sample passed the preservation cut-off and is well-preserved. The single sample that did not pass (from Farrer 2021) was excluded from all downstream analyses.

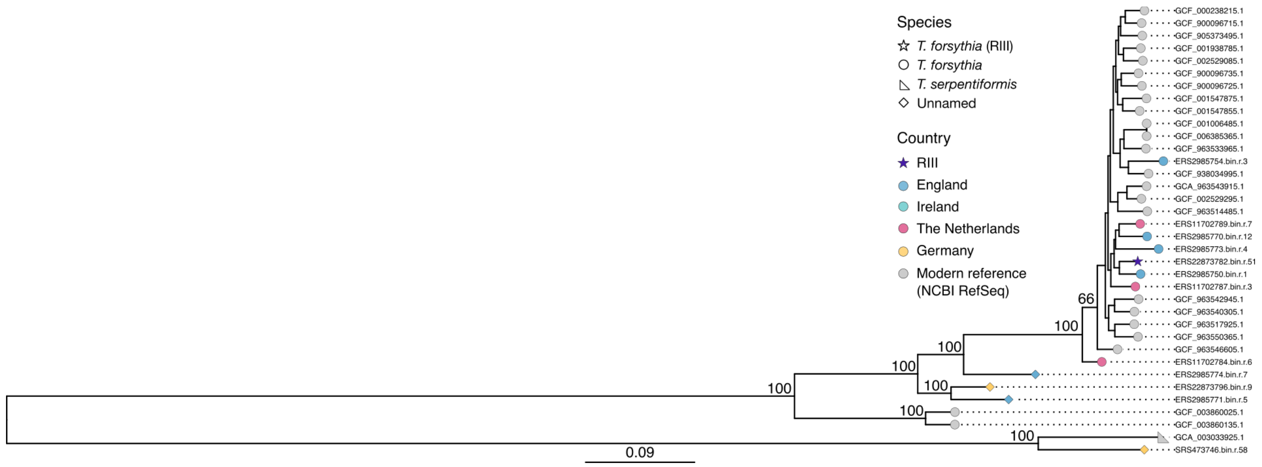


**Figure S2.** Tannerella forsythia genome phylogenetic tree with tree tips labeled by genome.

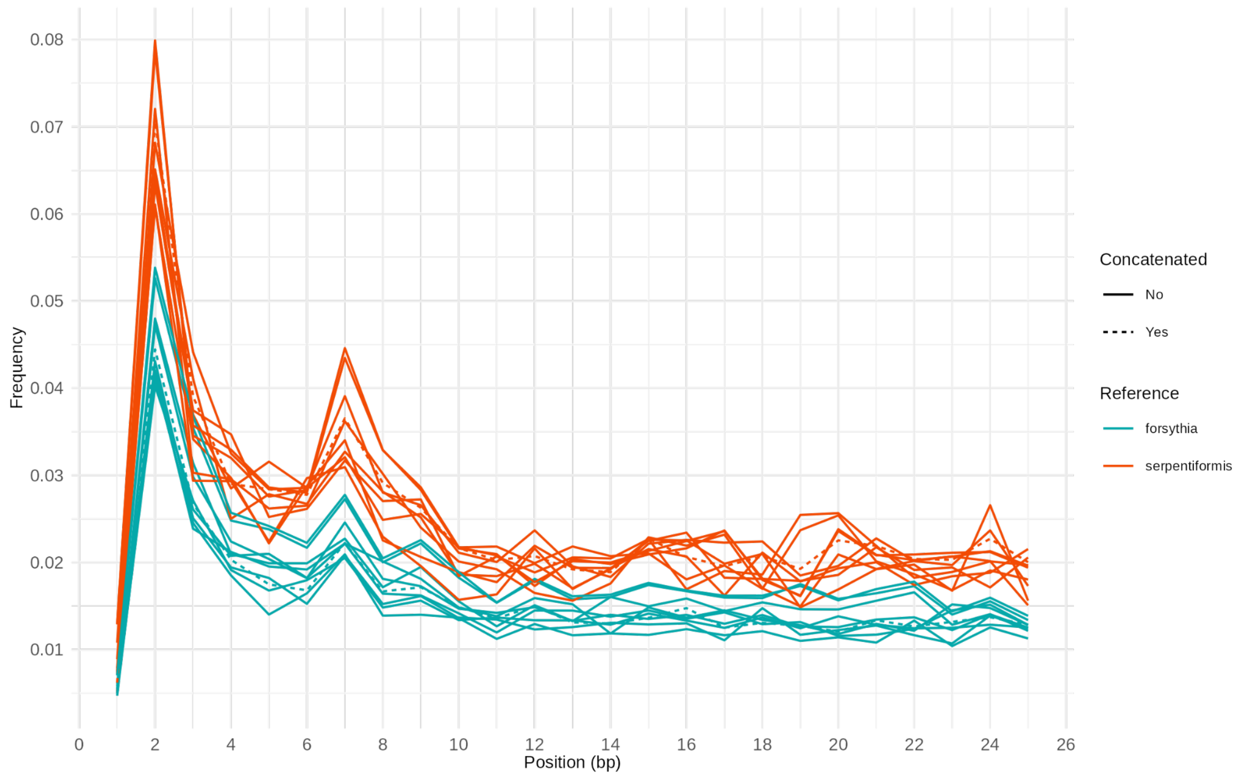


**Figure S3.** 5’-to-3’ C->T damage patterns on G12 libraries mapped against *Tannerella forsythia* or against *Tannerella serpentiformis*. Concatenated refers to all libraries, published and newly generated for this study, concatenated into a single fastq file for mapping. Non-concatenated indicates individual libraries. The libraries for sample G12 were built with the Phusion enzyme, which is a proofreading enzyme and therefore the measured damage is low.
